# A Federated and Privacy-Preserving Framework for Large-Scale Genome-Wide Association Studies with Mixed-Effects Models

**DOI:** 10.64898/2025.12.16.693409

**Authors:** Xiaowen Suo, Fuzhong Xue, Yanyan Zhao

## Abstract

Genome-wide association studies (GWAS) increasingly rely on large-scale data integration to achieve the statistical power necessary to detect variants with weak effects. However, genomic data are typically siloed across institutions, and privacy constraints often preclude centralized analysis. While federated learning (FL) offers a viable alternative by enabling cross-site computation without sharing individual-level data, applying mixed models, which are essential for correcting population structure, in a distributed setting remains a challenge in terms of statistical accuracy and computational scalability. Here, we present a federated mixed-model framework for GWAS that achieves high fidelity to centralized analyses while maintaining efficiency at biobank scale. Building on mixed-model theory and distributed optimization, we introduce algorithms for continuous (FedLMM) and binary (FedGLMM) traits that perform parameter estimation and association testing through site-local computation and aggregation of intermediate statistics. Comprehensive simulations spanning varied sample sizes and genomic densities demonstrate that our methods closely mirror centralized benchmarks (fastGWA and fastGWA-GLMM). Effect-size estimates exhibit near-perfect correlation, and over 99% of significant loci are recovered with well-controlled type I error rates. Empirical analyses on ∼100,000 UK Biobank participants further confirm that the framework delivers consistent inference while sustaining high computational performance. This work establishes a practical, open-source, and statistically reliable federated solution for large-scale GWAS, resolving the tension between data privacy and the need for statistical power in modern genomics.

## Introduction

The rapid expansion of genomic data has transformed biomedical research, enabling unprecedented opportunities to uncover the genetic architecture of complex traits and diseases^1^. Genome-wide association studies (GWAS) provide a systematic framework for linking genetic variants to phenotypic outcomes, revealing key loci involved in disease mechanisms and biological pathways^2,3^. However, the statistical power of GWAS fundamentally depends on two factors: the robustness of statistical models and the scale of sample sizes^4^. While methodological advances such as mixed-model approaches^5-12^ have improved control of population stratification and confounding, increasing sample size remains the most direct path to higher statistical power and broader generalizability. Yet, in practice, genomic datasets are fragmented across institutions, forming isolated “data silos”, and individual cohorts often lack sufficient power—especially for rare variants and population-specific effects.

Collaborative GWAS analysis across multiple institutions could overcome these limitations by aggregating data across diverse cohorts. However, direct data sharing is frequently infeasible due to legal, ethical, and technical barriers. Stringent regulations such as the General Data Protection Regulation (GDPR)^13^ and the Health Insurance Portability and Accountability Act (HIPAA)^14^ explicitly restrict the transfer of individual-level genetic data. Beyond regulatory concerns, centralized aggregation entails high operational costs, complex governance, and unresolved questions of data ownership. Large-scale initiatives (e.g., the UK Biobank^15^) illustrate the scientific value of massive integration but also the substantial financial and logistical burden required to achieve it. Consequently, valuable genomic resources in regional or institutional repositories remain underutilized, limiting the global diversity and representativeness of GWAS findings.

Federated learning (FL) has emerged as a compelling paradigm for collaborative analysis under data-locality constraints^16,17^. By allowing institutions to jointly optimize a shared model without exchanging raw data, FL enables privacy-preserving computation at scale. Nonetheless, applying FL to GWAS introduces distinctive computational and methodological hurdles, including the need to handle extremely high-dimensional genomic data, model random effects to account for genetic relatedness and population structure, and maintain statistical accuracy under strict data locality constraints. Existing federated GWAS frameworks^18-25^ often simplify the underlying models at the expense of statistical fidelity, or they rely on heavy cryptographic machinery that can limit practical scalability to biobank-scale analyses.

Here, we introduce a federated mixed-model framework designed to resolve these computational and privacy challenges. Our approach, encompassing FedLMM for continuous traits and FedGLMM for binary traits, performs all computations locally except for the exchange of aggregated intermediate parameters; no raw genotypes, phenotypes, or covariates leave institutional boundaries. By leveraging a three-stage optimization strategy, our framework drastically reduces the runtime of association testing without sacrificing statistical accuracy. We validate the system through comprehensive simulations and empirical analysis of ∼100,000 UK Biobank participants, demonstrating that it delivers results virtually identical to centralized benchmarks. This work establishes a scalable, privacy-preserving foundation for multi-cohort genomics, enabling secure collaboration that upholds data sovereignty while meeting the computational demands of large-scale genetic discovery.

## Results

### Overview of the Federated GWAS Framework

We developed a federated GWAS framework that supports large-scale mixed-model inference across multiple institutions without transferring individual-level data. Building on this architecture, we implemented two core algorithms—FedLMM for continuous traits and FedGLMM for binary traits—both designed for efficient distributed mixed-model GWAS.

The framework adopts a three-stage workflow that enables accurate and scalable inference: (1) **Initialization**: Fixed-effect parameters are rapidly initialized using distributed linear or logistic regression. (2) **Mixed Model Fitting**: Variance components are iteratively estimated via a distributed Penalized Quasi-Likelihood (PQL) approach combined with Average Information Restricted Maximum Likelihood (AI-REML). This step achieves precise parameter estimation without pooling individual-level data. (3) **Federated Score Testing**: Large-scale association testing is performed using a distributed GRAMMAR-Gamma approximation. By leveraging parallel computing, this stage achieves significant acceleration of the score test, overcoming the primary computational bottleneck of traditional mixed-model GWAS. An overview of the proposed federated learning framework is depicted in Fig. 1.

**Figure 1.**
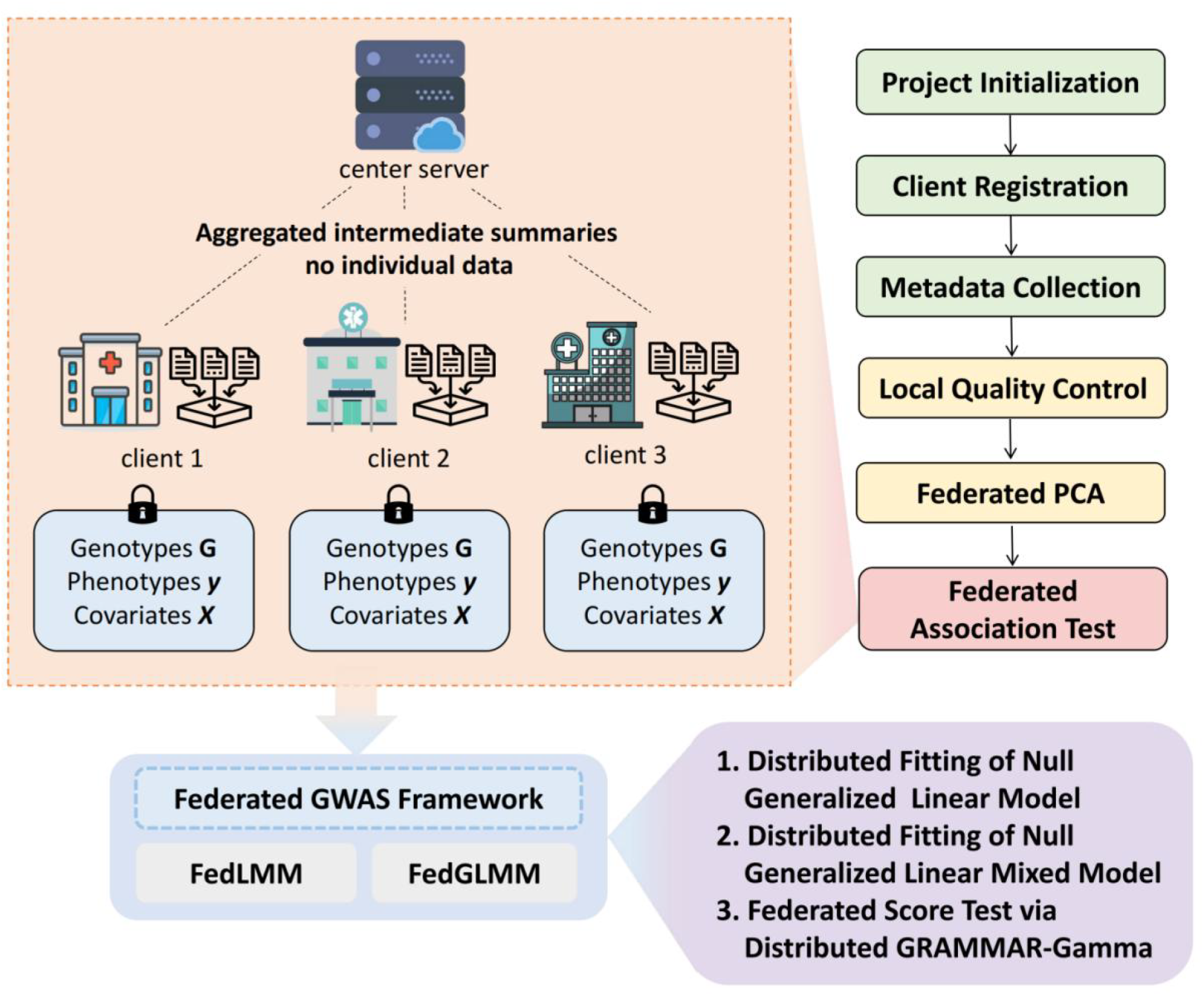
Privacy-preserving federated GWAS pipeline. Each site retains control over its raw genotypes, phenotypes, and covariates, sharing only aggregated intermediate summaries with a coordinating server. Sensitive computations are kept local throughout the process. The federated GWAS framework consists of the FedGLMM and FedLMM algorithms. FedLMM handles continuous traits, while FedGLMM is used for binary traits. The algorithms are organized into three stages: (1) initialization of fixed effects; (2) distributed mixed-model fitting using PQL and AI-REML updates; and (3) genome-wide association through a distributed GRAMMAR-Gamma score test.

The framework relies on a standard trust model where the central server coordinates aggregation protocols and clients faithfully execute local computations. Both algorithms are implemented in an open-source R package, providing a standardized and computationally efficient tool for secure multi-center genetic discovery.

### Concordance with Centralized Baselines across Simulation Settings

To evaluate the rigorousness of the proposed framework, we conducted extensive simulations varying sample sizes (6,000 to 15,000) and SNP densities (50,000, 100,000, 200,000) for both binary and continuous phenotypes. We benchmarked our federated algorithms against the centralized methods, fastGWA-GLMM and fastGWA (implemented in GCTA).

The fixed-effect coefficients estimated by our federated algorithms were virtually indistinguishable from the centralized estimates, with Pearson correlation coefficients consistently approaching 1.0. This confirms that our distributed optimization strategy successfully recovers global model parameters within numerical tolerance. Again, these results reflect one instance from the 100 simulations per scenario. Tab. 1 presents a comparison of the fixed-effect coefficients obtained by FedGLMM versus fastGWA-GLMM across 6 simulation scenarios, and Tab. 2 presents a similar comparison for FedLMM versus fastGWA across the same scenarios.

Quantile-quantile (Q-Q) plots of score statistics, variances, and P-values exhibited near-perfect alignment with those generated by centralized baselines. The test statistic distributions from the federated algorithms were virtually superimposable on the centralized results, demonstrating that our framework successfully replicates the statistical behavior of the centralized methods without requiring data centralization. These plots represent the results from one of the 100 simulations conducted per scenario. Fig. 3 presents the Q-Q plots of score statistics, variances, and P-values for simulation scenarios, comparing the results from the FedGLMM algorithm with those from the centralized fastGWA-GLMM benchmark. Fig. 5 presents similar Q-Q plots for the FedLMM algorithm, comparing its results with those from the centralized fastGWA benchmark.

**Figure 2.**
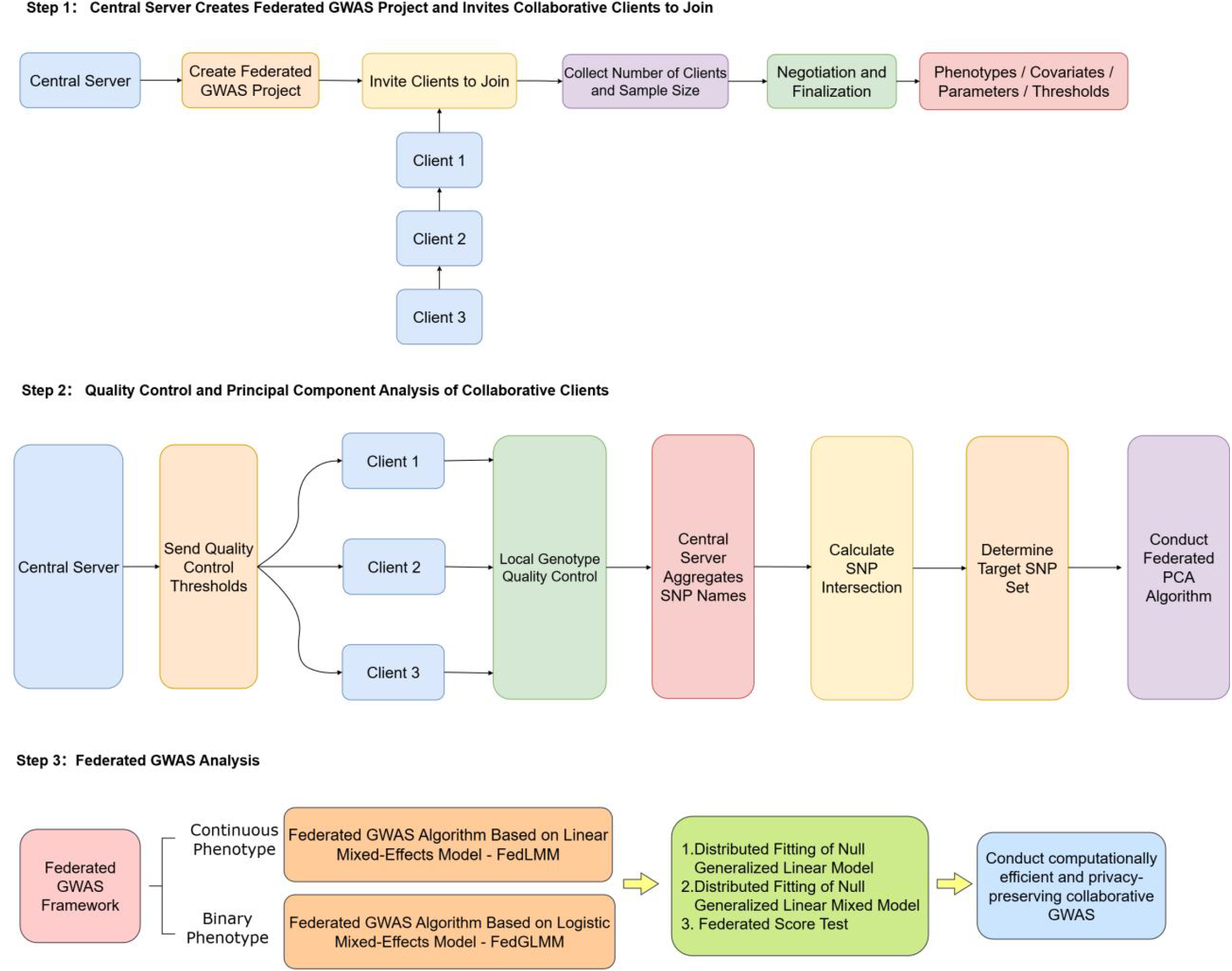
Overview of the Federated GWAS Workflow. This figure outlines the three main steps in the federated GWAS pipeline. Step 1: The central server initiates the project, invites clients, and finalizes information. Step 2: Quality control and PCA are performed, with SNP names aggregated by the central server to determine the target SNP set. Step 3: Federated GWAS analysis is conducted using FedLMM for continuous phenotypes and FedGLMM for binary phenotypes.

**Figure 3.**
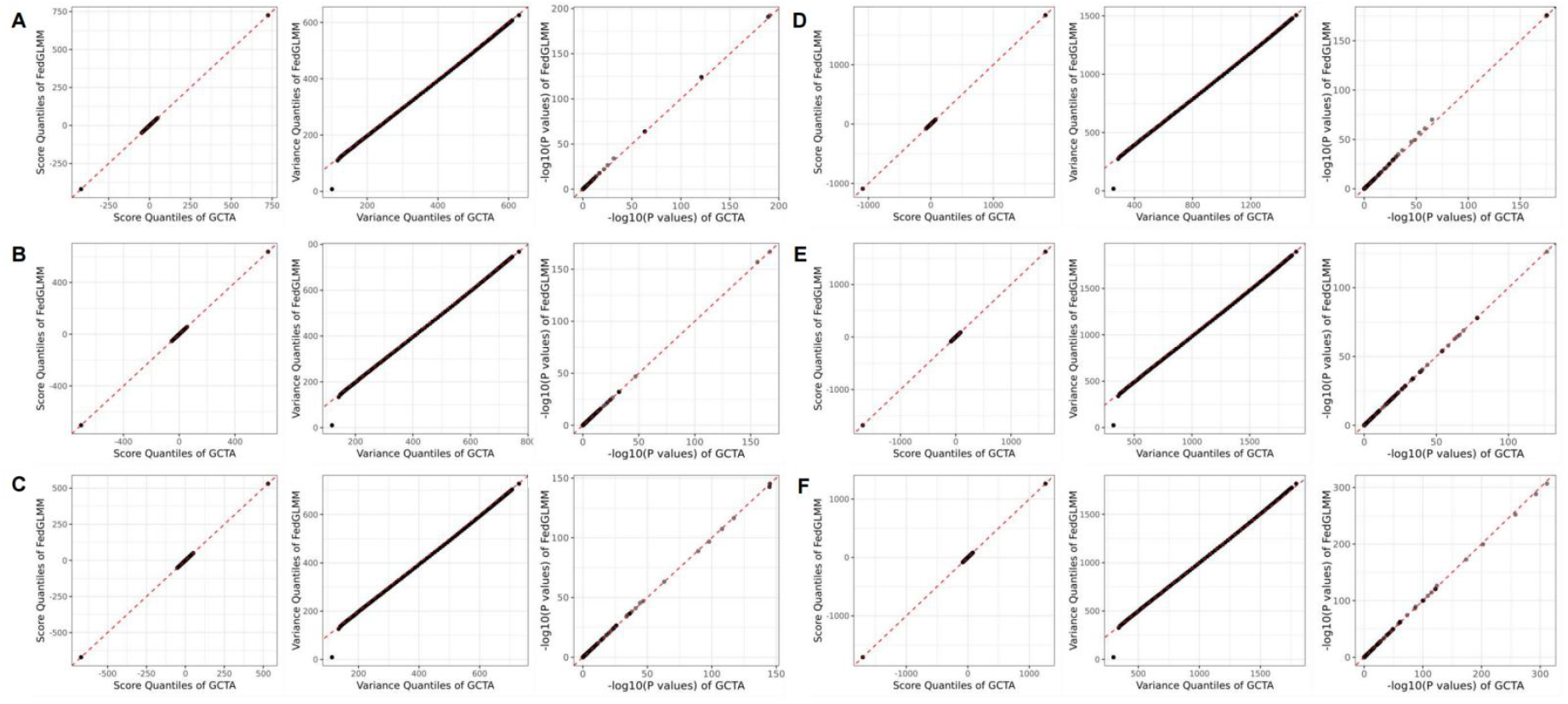
Q-Q plots of association metrics comparing FedGLMM and fastGWA-GLMM for simulation scenarios. Comparison of the performance of FedGLMM against GCTA across six simulation scenarios, varying in sample size (6,000 vs. 15,000 samples) and feature dimension (50,000, 100,000, 200,000 SNPs). (A) 6,000 samples, 50,000 SNPs; (B) 6,000 samples, 100,000 SNPs; (C) 6,000 samples, 200,000 SNPs; (D) 15,000 samples, 50,000 SNPs; (E) 15,000 samples, 100,000 SNPs; (F) 15,000 samples, 200,000 SNPs. Each plot compares the quantiles of the score, variance, and -log10(p-value) from FedGLMM (y-axis) with corresponding values from GCTA (x-axis). The red dashed line indicates perfect agreement between the two methods.

Circular Manhattan plots revealed near-perfect agreement in discovery. For binary phenotypes, the set of significant SNPs identified by FedGLMM overlapped >99.9% with those from fastGWA-GLMM. For contincious phenotypes, FedLMM achieved >99.0% concordance for continuous traits. These results are also derived from one of the 100 simulations performed for each scenario. Fig. 4 displays circular Manhattan plots for simulation scenarios, contrasting the association results obtained by the FedGLMM algorithm against the centralized fastGWA-GLMM benchmark. Fig. 6 displays the corresponding plots for FedLMM versus fastGWA.

**Figure 4.**
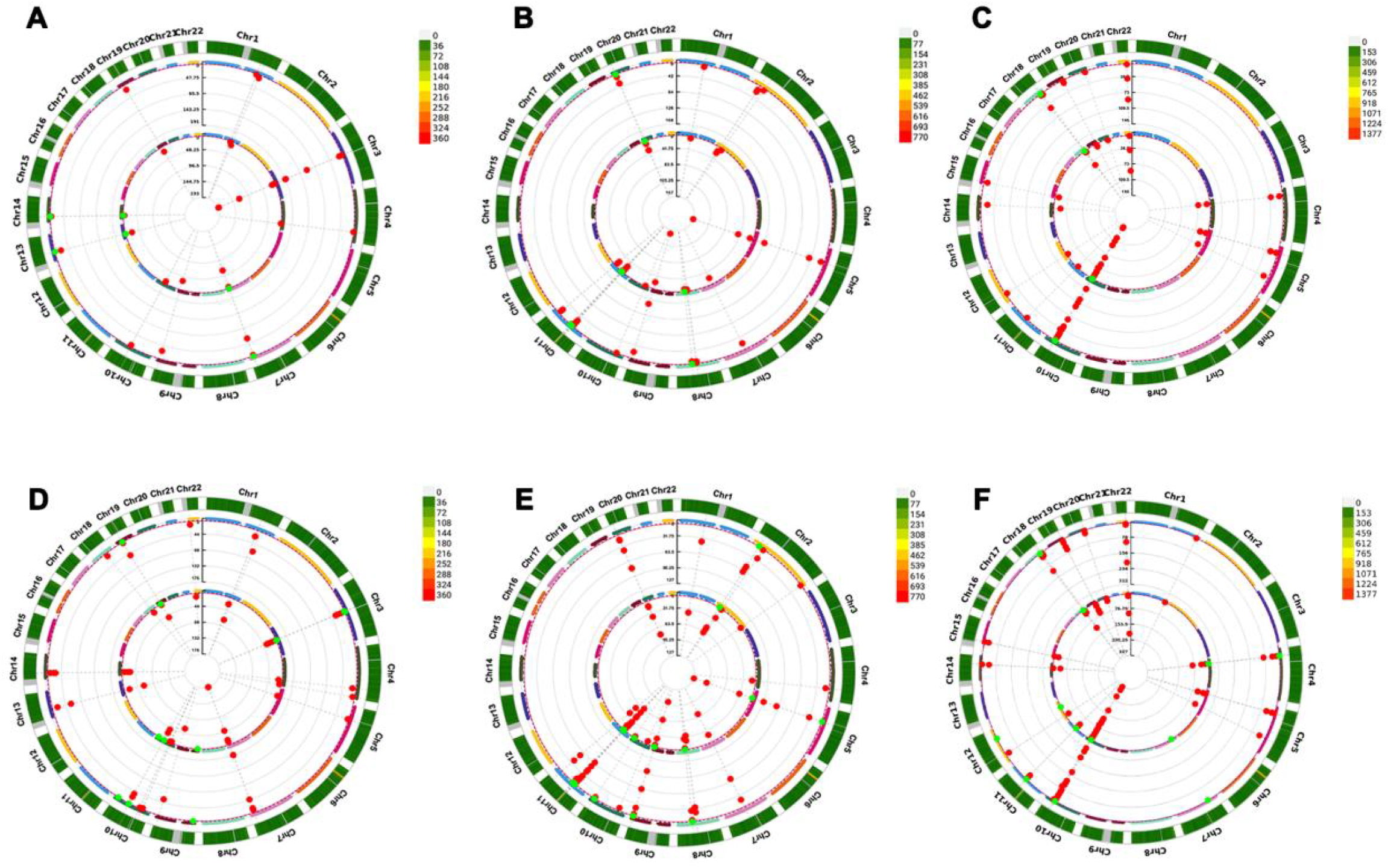
Circular Manhattan plots of association results comparing FedGLMM and fastGWA-GLMM for simulation scenarios. Genome-wide association results from FedGLMM closely match those from the centralized fastGWA-GLMM benchmark, confirming accurate signal detection. (A) 6,000 samples, 50,000 SNPs; (B) 6,000 samples, 100,000 SNPs; (C) 6,000 samples, 200,000 SNPs; (D) 15,000 samples, 50,000 SNPs; (E) 15,000 samples, 100,000 SNPs; (F) 15,000 samples, 200,000 SNPs.

It is important to note that in the comparison between FedLMM and fastGWA, only in the first simulation scenario, where the sample size was 6,000 and the number of SNPs was 50,000, did the statistics not match almost perfectly. As shown in the Fig. 5, the variance data points tightly cluster around the theoretical expected line, with near-perfect alignment. However, the effect estimates exhibited some divergence: a portion of the GCTA effect estimates were significantly smaller than those from the FedLMM algorithm. This difference in effect estimates subsequently led to variations in the p-values. Despite this discrepancy, both algorithms identified a similar number of significant SNPs, as the associated SNPs were pre-known in the simulation. However, GCTA identified significantly more significant SNPs than FedLMM, indicating that, while the power was comparable, GCTA had a higher false positive rate.

**Figure 5.**
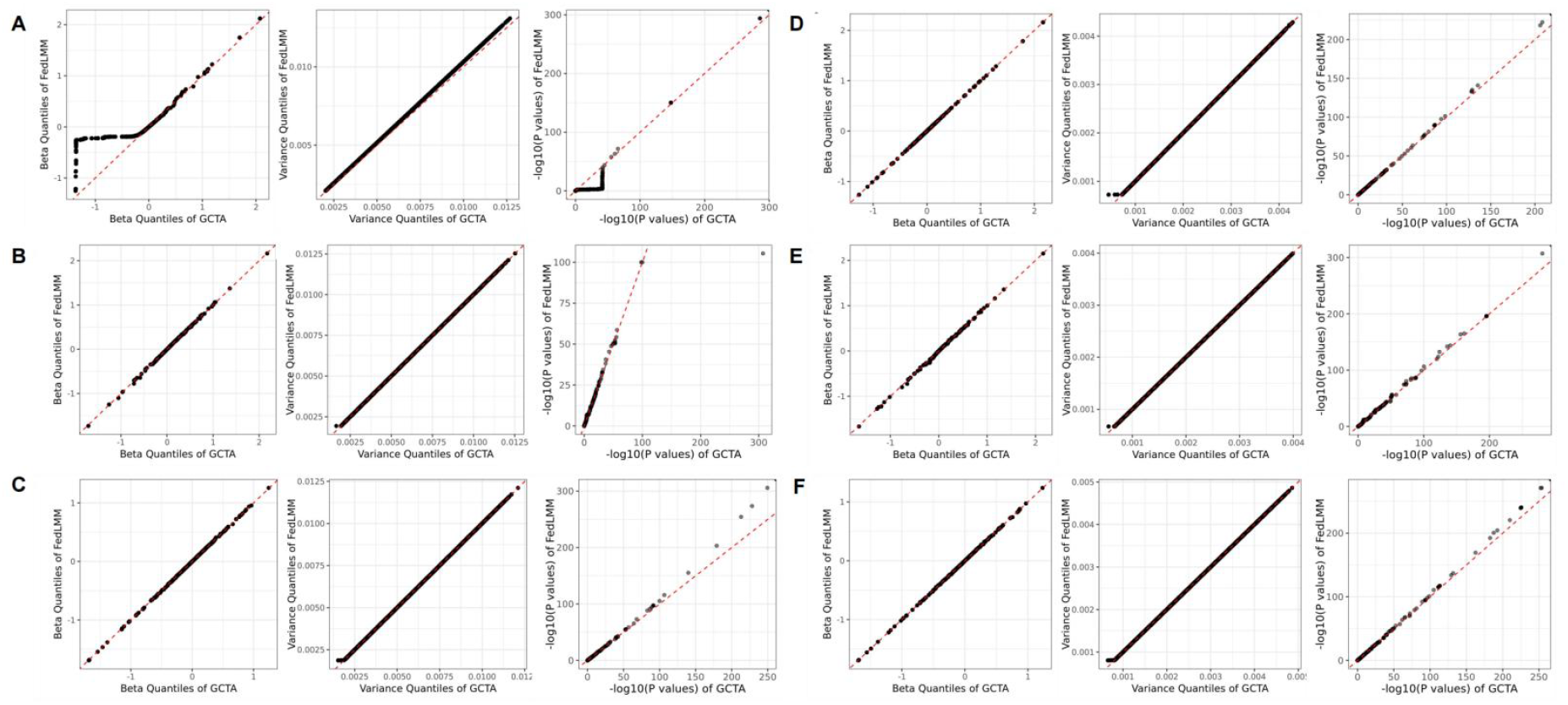
Q-Q plots of association metrics comparing FedLMM and fastGWA for simulation scenarios. Comparison of the performance of FedLMM against GCTA across six simulation scenarios, varying in sample size (6,000 vs. 15,000 samples) and feature dimension (50,000, 100,000, 200,000 SNPs). (A) 6,000 samples, 50,000 SNPs; (B) 6,000 samples, 100,000 SNPs; (C) 6,000 samples, 200,000 SNPs; (D) 15,000 samples, 50,000 SNPs; (E) 15,000 samples, 100,000 SNPs; (F) 15,000 samples, 200,000 SNPs. Each plot compares the quantiles of the score, variance, and -log10(p-value) from FedLMM (y-axis) with corresponding values from GCTA (x-axis). The red dashed line indicates perfect agreement between the two methods.

**Figure 6.**
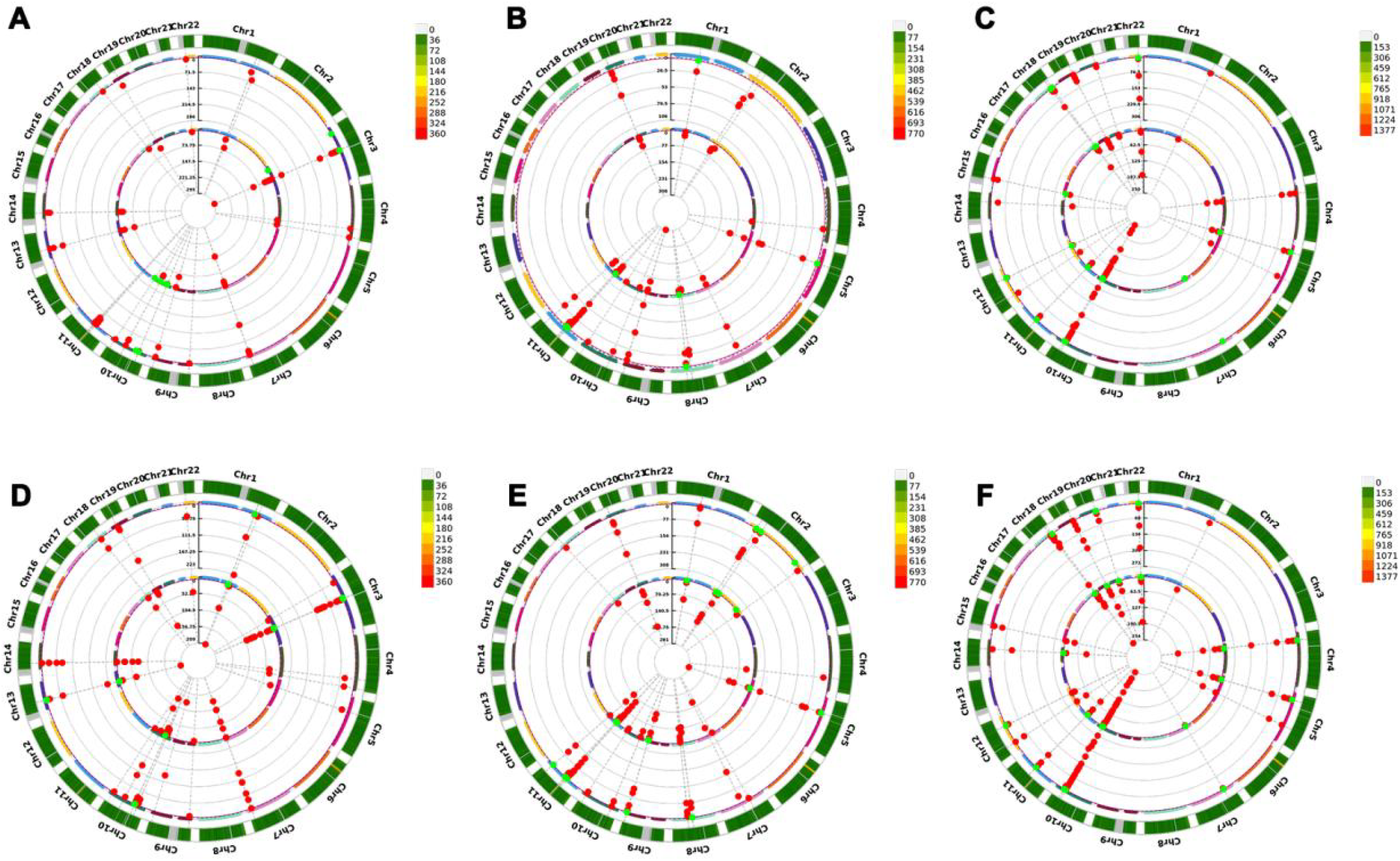
Circular Manhattan plots of association results comparing FedLMM and fastGWA for simulation scenarios. Genome-wide association results from FedLMM closely match those from the centralized fastGWA benchmark, confirming accurate signal detection. (A) 6,000 samples, 50,000 SNPs; (B) 6,000 samples, 100,000 SNPs; (C) 6,000 samples, 200,000 SNPs; (D) 15,000 samples, 50,000 SNPs; (E) 15,000 samples, 100,000 SNPs; (F) 15,000 samples, 200,000 SNPs.

Beyond estimation accuracy, we assessed the inferential reliability of the framework by quantifying statistical power and False Positive Rates (FPR). FedLMM demonstrated a highly favorable trade-off between sensitivity and specificity. It achieved a detection power of 0.841, comparable to the centralized baseline (0.850). Notably, FedLMM exhibited superior control of false positives, with an FPR of 0.00114—substantially lower than that of fastGWA (0.00265). This suggests that our distributed implementation maintains rigorous Type I error control, effectively minimizing spurious findings while retaining high sensitivity to true genetic signals. FedGLMM achieved inferential performance identical to the centralized fastGWA-GLMM. Both methods yielded an identical power of 0.708 and an FPR of 0.000925. This alignment confirms that the distributed strategy employed in FedGLMM effectively preserves the global statistical properties, achieving inferential equivalence to the centralized baseline.

Collectively, these simulations validate that the proposed framework delivers empirical equivalence to centralized GWAS in terms of test statistic distribution, discovery overlap variants, and fixed-effect estimation, while offering robust discovery capabilities and strict adherence to significance thresholds in centralized settings.

### Empirical Validation Using the UK Biobank

To demonstrate the practical utility of our framework, we applied FedLMM and FedGLMM to real-world data from the UK Biobank. The analysis included 97,007 samples partitioned into three geographic cohorts: Northern (44,253), Central (23,597), and Southern England (29,157). After harmonizing genotype data across sites—ensuring non-missing data comparability for all participants—a final set of 296,961 common variants was retained for association testing.

The results were effectively indistinguishable from those obtained by the centralized “gold standard” tools, fastGWA and fastGWA-GLMM. For both BMI (continuous) and smoking status (binary), the federated algorithms yielded fixed-effect coefficients and score test statistics that were nearly identical to centralized estimates. The p-value distributions showed rigorous concordance, confirming that the distributed optimization strategy successfully reconstructs the global model results without data centralization. The comparison of fixed-effect coefficients for both binary and continuous phenotypes across the federated and centralized methods is provided in the Tab. 6.

Using a genome-wide significance threshold of 5×10^-8^, FedGLMM identified seven significant loci, fully replicating the signals detected by fastGWA-GLMM. Notable associations included rs1447481, a well-established risk variant for smoking behavior, rs4547132, previously linked to educational attainment (a known correlate of smoking), and rs4144892, associated with smoking initiation. The remaining four hits were rare variants, highlighting the framework’s sensitivity across the allele frequency spectrum. For BMI, FedLMM identified 161 significant associations, showing substantial overlap (158 shared SNPs) with the 164 hits found by GCTA. The minor discrepancy in the number of marginally significant hits is attributable to slight numerical differences in the iterative optimization paths, yet the core biological signals remained robust.

### Computational Performance

We systematically evaluated the computational performance of FedGLMM and FedLMM in terms of runtime, memory usage, and communication cost across varying data scales. In our simulated federated environment (three clients and one central server), both algorithms demonstrated linear scalability with respect to sample size and SNP count, confirming their feasibility for biobank-scale analyses.

In terms of runtime (Table 3), both algorithms demonstrated predictable scalability. FedLMM exhibited exceptional speed, completing the analysis of 15,000 samples and 200,000 SNPs in just about 3.9 minutes (233.8 s), with its runtime scaling approximately linearly with SNP count. FedGLMM, tasked with solving the more computationally intensive generalized mixed model, required about 57 minutes (3426.8 s) for the same dataset. Notably, the bulk of the computational cost for FedGLMM lies in the score test phase, which scales linearly with the number of variants, whereas the null model fitting remains relatively efficient and stable.

**Table 1.**
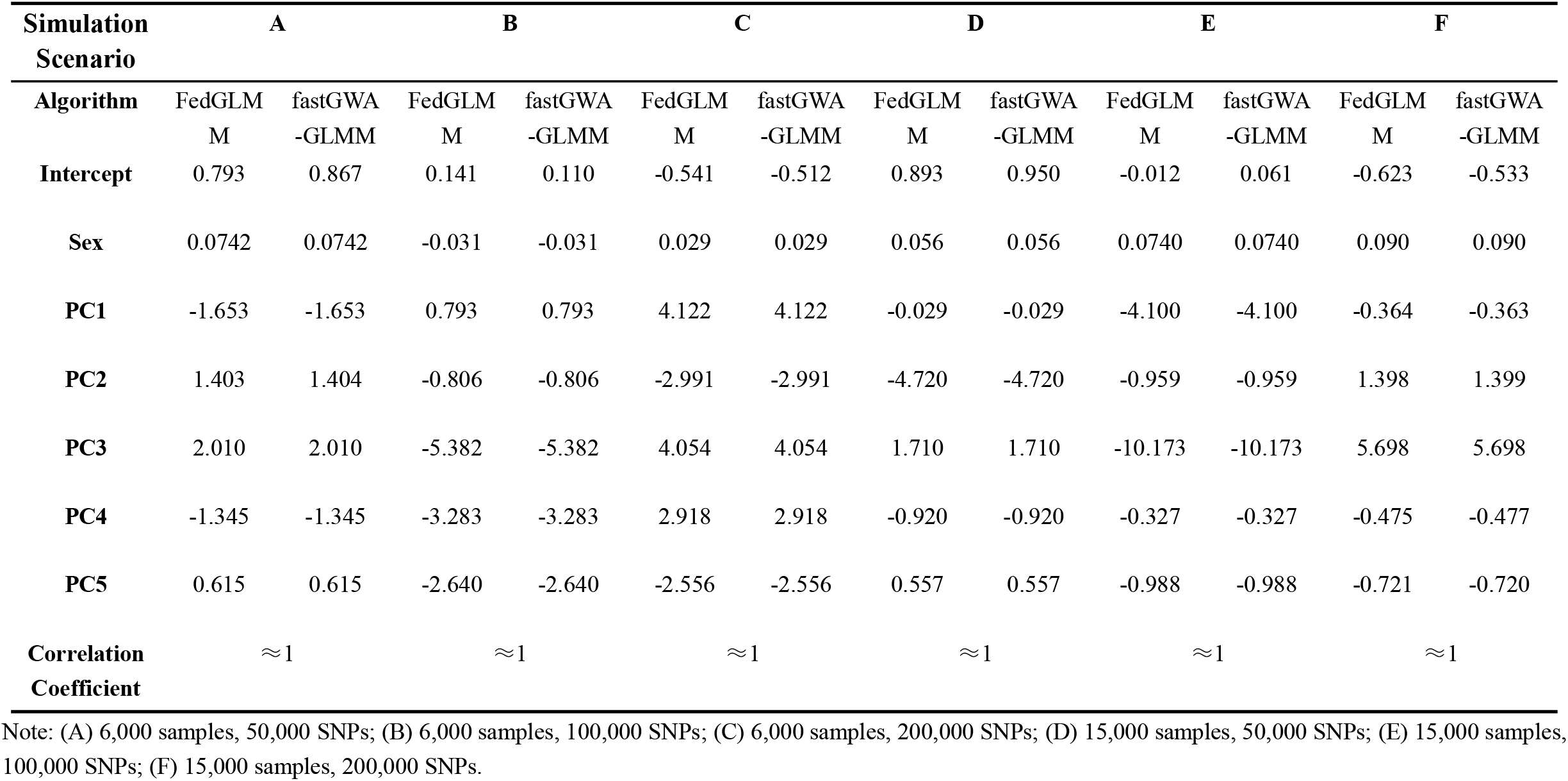
Comparison of Fixed Effect Coefficients Between FedGLMM and fastGWA-GLMM Across Simulation Scenarios.

| **Simulation Scenario** | **A** |  | **B** |  | **C** |  | **D** |  | **E** |  | **F** |  |
| --- | --- | --- | --- | --- | --- | --- | --- | --- | --- | --- | --- | --- |
| **Algorithm** | FedGLM<br>M | fastGWA<br>-GLMM | FedGLM<br>M | fastGWA<br>-GLMM | FedGLM<br>M | fastGWA<br>-GLMM | FedGLM<br>M | fastGWA<br>-GLMM | FedGLM<br>M | fastGWA<br>-GLMM | FedGLM<br>M | fastGWA<br>-GLMM |
| **Intercept** | 0.793 | 0.867 | 0.141 | 0.110 | -0.541 | -0.512 | 0.893 | 0.950 | -0.012 | 0.061 | -0.623 | -0.533 |
| **Sex** | 0.0742 | 0.0742 | -0.031 | -0.031 | 0.029 | 0.029 | 0.056 | 0.056 | 0.0740 | 0.0740 | 0.090 | 0.090 |
| **PC1** | -1.653 | -1.653 | 0.793 | 0.793 | 4.122 | 4.122 | -0.029 | -0.029 | -4.100 | -4.100 | -0.364 | -0.363 |
| **PC2** | 1.403 | 1.404 | -0.806 | -0.806 | -2.991 | -2.991 | -4.720 | -4.720 | -0.959 | -0.959 | 1.398 | 1.399 |
| **PC3** | 2.010 | 2.010 | -5.382 | -5.382 | 4.054 | 4.054 | 1.710 | 1.710 | -10.173 | -10.173 | 5.698 | 5.698 |
| **PC4** | -1.345 | -1.345 | -3.283 | -3.283 | 2.918 | 2.918 | -0.920 | -0.920 | -0.327 | -0.327 | -0.475 | -0.477 |
| **PC5** | 0.615 | 0.615 | -2.640 | -2.640 | -2.556 | -2.556 | 0.557 | 0.557 | -0.988 | -0.988 | -0.721 | -0.720 |
| **Correlation Coefficient** | $\approx 1$ | | $\approx 1$ | | $\approx 1$ | | $\approx 1$ | | $\approx 1$ | | $\approx 1$ | |

Note: (A) 6,000 samples, 50,000 SNPs; (B) 6,000 samples, 100,000 SNPs; (C) 6,000 samples, 200,000 SNPs; (D) 15,000 samples, 50,000 SNPs; (E) 15,000 samples, 100,000 SNPs; (F) 15,000 samples, 200,000 SNPs.

**Table 2.** Comparison of Fixed Effect Coefficients Between FedLMM and fastGWA Across Simulation Scenarios.

| **Simulation Scenario** | **A** |  | **B** |  | **C** |  | **D** |  | **E** |  | **F** |  |
| --- | --- | --- | --- | --- | --- | --- | --- | --- | --- | --- | --- | --- |
| **Algorithm** | FedLMM | fastGWA | FedLMM | fastGWA | FedLMM | fastGWA | FedLMM | fastGWA | FedLMM | fastGWA | FedLMM | fastGWA |
| **Intercept** | 1.570 | 1.570 | 0.108 | 0.109 | -1.093 | -1.093 | 1.696 | 1.696 | 0.041 | 0.041 | -1.135 | -1.135 |
| **Sex** | 0.092 | 0.092 | 0.081 | 0.081 | 0.155 | 0.155 | 0.064 | 0.064 | 0.112 | 0.112 | 0.180 | 0.180 |
| **PC1** | 0.222 | 0.222 | 0.738 | 0.738 | 2.729 | 2.730 | -1.367 | -1.365 | -5.356 | -5.355 | -3.025 | -3.024 |
| **PC2** | 0.795 | 0.795 | 1.388 | 1.387 | -1.861 | -1.861 | -4.141 | -4.141 | -3.149 | -3.149 | 0.252 | 0.252 |
| **PC3** | -2.336 | -2.336 | -13.455 | -13.455 | 7.594 | 7.594 | 3.723 | 3.723 | -18.980 | -18.980 | 10.409 | 10.408 |
| **PC4** | -0.983 | -0.983 | -2.096 | -2.096 | 1.533 | 1.533 | -1.719 | -1.719 | 1.325 | 1.325 | 0.146 | 0.146 |
| **PC5** | -0.992 | -0.992 | -5.453 | -5.453 | -0.141 | -0.141 | 4.243 | 4.243 | -1.018 | -1.018 | 0.017 | 0.017 |
| **Correlation Coefficient** | $\approx 1$ | | $\approx 1$ | | $\approx 1$ | | $\approx 1$ | | $\approx 1$ | | $\approx 1$ | |

Note: (A) 6,000 samples, 50,000 SNPs; (B) 6,000 samples, 100,000 SNPs; (C) 6,000 samples, 200,000 SNPs; (D) 15,000 samples, 50,000 SNPs; (E) 15,000 samples, 100,000 SNPs; (F) 15,000 samples, 200,000 SNPs.

**Table 3.** Summary table of algorithm runtime in simulation scenarios.

| Algorithm | Sample Size &<br>SNP Count | Step 1 Runtime<br>(s) | Step 2 Runtime<br>(s) | Total<br>Runtime (s) |
| --- | --- | --- | --- | --- |
| **FedGLMM** | 6000,50000 | 61.0 | 86.5 | 147.5 |
|  | 6000,100000 | 62.3 | 139.2 | 201.5 |
|  | 6000,200000 | 63.6 | 268.1 | 331.7 |
|  | 15000,50000 | 134.0 | 834.8 | 968.8 |
|  | 15000,100000 | 123.2 | 1033.7 | 1156.9 |
|  | 15000,200000 | 86.2 | 3340.6 | 3426.8 |
| **FedLMM** | 6000,50000 | 14.4 | 24.9 | 39.3 |
|  | 6000,100000 | 13.6 | 40.9 | 54.5 |
|  | 6000,200000 | 13.6 | 68.9 | 82.5 |
|  | 15000,50000 | 59.0 | 63.9 | 122.9 |
|  | 15000,100000 | 61.2 | 107.9 | 169.1 |
|  | 15000,200000 | 61.7 | 172.1 | 233.8 |

Note: All time values in this table are reported in seconds (s). The time for Step 1 in this table includes the total time for fitting the generalized linear model and the generalized linear mixed-effects model.

Regarding memory usage (Table 4), consumption remained highly manageable. For the largest simulation case (N=15,000, p=200,000), the cumulative memory for all three clients and the server was only 16.3 GB for FedGLMM and 18.8 GB for FedLMM. This indicates that the per-site memory requirement is relatively low, making the framework deployable even on resource-constrained hardware. Memory usage showed a stronger dependence on SNP count during the association testing step than on sample size, consistent with the storage requirements for genotype matrices and summary statistics.

**Table 4.** Summary table of algorithm running memory in simulation scenarios.

| Algorithm | Sample Size &<br>SNP Count | Step 1 Memory<br>(GB) | Step 2 Memory<br>(GB) | Total Memory<br>(GB) |
| --- | --- | --- | --- | --- |
| **FedGLMM** | 6000,50000 | 0.5 | 1.4 | 1.9 |
|  | 6000,100000 | 0.6 | 2.5 | 3.1 |
|  | 6000,200000 | 0.6 | 4.7 | 5.3 |
|  | 15000,50000 | 3.5 | 4.4 | 7.9 |
|  | 15000,100000 | 3.5 | 7.2 | 10.7 |
|  | 15000,200000 | 3.5 | 12.8 | 16.3 |
| **FedLMM** | 6000,50000 | 0.5 | 1.8 | 2.4 |
|  | 6000,100000 | 0.6 | 3.0 | 3.6 |
|  | 6000,200000 | 0.6 | 5.2 | 5.8 |
|  | 15000,50000 | 3.4 | 6.9 | 10.3 |
|  | 15000,100000 | 3.2 | 9.7 | 13.1 |
|  | 15000,200000 | 3.3 | 15.3 | 18.8 |

Note: All memory values in this table are reported in gigabytes (GB). The memory for Step 1 in this table includes the total memory usage for fitting the generalized linear model and the generalized linear mixed-effects model.

Communication overhead, a critical bottleneck in federated learning, is summarized in Table 5. Our theoretical analysis confirms that the data transfer volume is determined by algorithmic parameters rather than raw data size. Specifically, in Step 2 (Mixed Model Fitting), communication cost is proportional to the number of sites and the square of the covariate count but is independent of the total sample size, ensuring that bandwidth requirements do not explode as biobanks grow larger. In Step 3 (Association Testing), communication is proportional to the number of SNPs as aggregated score statistics are exchanged. However, since no individual-level data is transmitted, the total transfer volume remains a small fraction of the raw dataset size, ensuring bandwidth efficiency.

**Table 5.** Summary table of communication costs between clients and central servers.

| Algorithm | Steps | Client -> Server<br>Transmission | Server -> Client<br>Transmission |
| --- | --- | --- | --- |
| **FedGLMM** | Step 1 - Null model Fitting | $\tilde{\ell}_j, \mathbf{H}_j$ | / |
| | Step 2 - Null mixed model Fitting | $\Sigma_j^{-1} \mathbf{X}_j, \mathbf{X}_j^T \Sigma_j^{-1} \mathbf{X}_j$ | $\alpha, \Sigma_j^{-1}$ |
| | Step 3 -Federated Score test | $\mathbf{G}_j^T \mathbf{res}_j, \mathbf{Z}_j \mathbf{G}_j, \tilde{\mathbf{G}}_j$ | $\mathbf{res}_j, \mathbf{Z}_j \mathbf{G}_j$ |
| **FedLMM** | Step 1 - Null model Fitting | $\tilde{\ell}_j, \mathbf{H}_j$ | / |
| | Step 2 - Null mixed model Fitting | $\Sigma_j^{-1} \mathbf{X}_j, \mathbf{X}_j^T \Sigma_j^{-1} \mathbf{X}_j$ | $\alpha, \Sigma_j^{-1}$ |
| | Step 3 -Federated Score test | $\dot{\mathbf{G}}_j^T \mathbf{V}_j^{-1} \dot{\mathbf{G}}_j, \dot{\mathbf{G}}_j^T \dot{\mathbf{G}}_j$ | / |

**Table 6.** Comparison of Fixed Effect Coefficients Between FedLMM and fastGWA in Empirical Analysis.

|  | FedGLMM | fastGWA-GLMM | FedLMM | fastGWA |
| --- | --- | --- | --- | --- |
| **Intercept** | 0.610 | 0.611 | -0.992 | -0.991 |
| **Sex** | -4.786 | -4.857 | 0.291 | 0.291 |
| **Age** | -1.054 | -1.088 | 1.951 | 1.950 |
| **Sex\*Age** | 13.082 | 13.259 | 2.200 | 2.200 |
| **Age\*Age** | 0.668 | 0.701 | -1.547 | -1.547 |
| **Sex\*Age<sup>2</sup>** | -9.321 | -9.431 | -1.842 | -1.842 |
| **PC1** | 33.713 | 33.716 | 1.274 | 1.274 |
| **PC2** | 0.517 | 0.518 | -2.609 | -2.609 |
| **PC3** | -2.203 | -2.203 | -0.357 | -0.357 |
| **PC4** | -6.695 | -6.696 | -1.819 | -1.820 |
| **PC5** | 13.246 | 13.246 | 0.146 | 0.146 |
| **PC6** | 7.771 | 7.772 | -1.500 | -1.500 |
| **PC7** | -9.146 | -9.146 | 1.169 | 1.170 |
| **PC8** | 5.312 | 5.313 | -2.325 | -2.324 |
| **PC9** | 10.137 | 10.136 | -0.903 | -0.903 |
| **PC10** | -0.618 | -0.618 | -0.098 | -0.097 |

To validate performance at a biobank scale, we conducted an empirical analysis using approximately 100,000 samples from the UK Biobank. FedLMM maintained high efficiency, completing the entire pipeline for BMI (continuous trait) in approximately 1.2 hours, with a total memory usage of around 203 GB across all simulated nodes. FedGLMM required a higher computational load for smoking status (binary trait). The pipeline took approximately 57.5 hours in total (Step 1: ∼1.25 hours, Step 2: ∼56.3 hours), with a total memory usage of approximately 210 GB across all simulated nodes. It is important to note that the reported memory usage represents the aggregate consumption of four logical nodes (three clients and one server); thus, the actual per-site memory requirement is lower and well within the capabilities of standard institutional servers.

Finally, compared to existing federated GLMM frameworks such as FedGMMAT, our algorithms significantly accelerated the association testing phase, improving computational efficiency. Specifically, while FedGMMAT encounters severe memory and computational bottlenecks when processing datasets with about 100,000 individuals, often resulting in memory overflow and failure to complete the analysis, our algorithm effectively handles such large-scale datasets without such issues. This optimization enabled faster processing times, particularly in large-scale datasets, by reducing the computational limitations commonly encountered during association testing.

## Discussion

This study presents a federated learning framework for GWAS that achieves both scalability and privacy preservation without compromising analytical precision. Through the development of FedLMM and FedGLMM, we demonstrate that large-scale mixed-model GWAS can be effectively conducted across multiple institutions while maintaining near-identical accuracy to centralized methods. The proposed framework integrates distributed PQL, AI-REML, and a novel distributed parallelized GRAMMAR-Gamma approximation to enable computationally efficient and statistically robust association testing under strict data locality constraints. Our results, validated through both simulation and empirical analysis on the UK Biobank dataset, highlight the framework’s ability to deliver high performance, reproducibility, and robustness. The proposed framework establishes a new computational paradigm for collaborative genetic research in the era of distributed data governance.

Unlike existing privacy-preserving and federated GWAS methods that rely heavily on cryptographic techniques or meta-analysis approximations, our framework adopts an algorithmic design that avoids raw data transmission by construction. By transmitting only aggregated intermediate parameters, we eliminate the need for encryption-based computation, which is often associated with high latency and memory costs. This design choice allows the framework to scale effectively to biobank-level datasets, achieving computational performance on par with centralized analysis.

To rigorously evaluate the accuracy of our federated algorithms, we benchmarked them against GCTA (specifically fastGWA and fastGWA-GLMM), a widely recognized centralized toolkit for large-scale mixed-model GWAS. Although recent methods like REGENIE offer computational advantages for biobank-scale data, their two-step regression strategy—fitting a null model to generate polygenic predictors as covariates—fundamentally differs from the joint mixed-model approach employed in our framework. As a result, fastGWA and fastGWA-GLMM were selected as the most theoretically aligned “gold standard” to ensure a fair comparison.

Current federated GWAS methods based on mixed-effect models can be categorized into two main approaches: the SafeGENIE/SF-GWAS framework and FedGMMAT. The SafeGENIE^26^ and SF-GWAS^27^ frameworks, developed by Hyunghoon Cho et al., integrate multi-party homomorphic encryption with distributed linear regression models and use stacked ridge regression for linear mixed models. These methods provide a secure and systematic solution, with SF-GWAS offering an expanded workflow that includes PCA. The key advantage of this approach lies in its robust cryptographic security, making it suitable for high-security international collaborations. The sfkit toolkit, built on this framework, translates these complex methods into practical tools, significantly lowering the barrier for federated GWAS applications^28^. Its main contribution is the integration of cryptographic protocols with bioinformatics workflows, enabling biologists to apply advanced security techniques directly. In contrast, the FedGMMAT method, proposed by Li et al., is based on generalized linear mixed models and corrects for both fixed and random effects across sites^29^. It also integrates homomorphic encryption for intermediate statistics. Although it shows promise in datasets with thousands of samples, it encounters significant computational bottlenecks when scaled to larger datasets, especially when sample sizes exceed tens of thousands or when working with large SNP sets. In such cases, the time and memory requirements for a full analysis exceed acceptable limits, highlighting the need for further optimization in large-scale genomic studies.

Despite these advances, several limitations remain. First, the current algorithms assume sample independence, which may restrict their application to family-based or hierarchically structured data. Extending the model to incorporate genetic relatedness matrices (GRMs) would enhance its applicability but also increase computational cost. Second, while the implemented privacy mechanism effectively prevents direct data leakage, it does not yet incorporate formal cryptographic protections such as homomorphic encryption^30,31^ or differential privacy^32^. Future work will explore integrating such lightweight encryption methods to further strengthen resistance to adversarial inference attacks while maintaining computational feasibility. Third, communication efficiency remains a bottleneck in large-scale deployments. The current iterative global–local optimization requires multiple communication rounds between clients and the central server. As the number of participating sites or the data dimensionality grows, communication delays and bandwidth consumption may increase substantially. Incorporating communication compression or asynchronous aggregation strategies could significantly mitigate these issues.

Importantly, the distributed optimization principles developed in this study have strong generalizability. Beyond GWAS, the proposed framework can be readily extended to other biomedical data analysis scenarios requiring secure multi-center modeling. For example, within electronic health record systems, it could be used to build distributed generalized mixed models for disease risk prediction, treatment effect evaluation, and pharmacovigilance across multiple hospitals while ensuring strict patient privacy. This adaptability underscores the potential of the framework to serve as a methodological foundation for privacy-preserving collaborative analysis in precision medicine.

In summary, this study establishes a high-fidelity, computationally scalable, and privacy-preserving federated framework for mixed-model GWAS. By demonstrating that biobank-scale association analyses can be conducted collaboratively without raw data sharing, it provides a methodological foundation for next-generation federated genomic analytics. The integration of distributed optimization, mixed-model inference, and privacy-aware system design offers a principled computational approach that can be broadly extended to secure scientific collaboration across heterogeneous, privacy-constrained environments.

## Methods

### 1 Federated GWAS Workflow

To ensure privacy preservation, raw genotype, covariate, and phenotype data are kept entirely local and never exchanged between participating clients and the central server. The federated GWAS workflow, illustrated in Fig. 2, proceeds through three primary stages:

1. **Project Initialization and Consensus Building**: The central server initiates the project by inviting participating clients and collecting metadata, including the number of clients and their sample sizes. Key parameters such as the target phenotype, covariates, quality control thresholds, number of principal components (PCs), and algorithm settings are defined by the central server, ensuring a consensus across all sites. This initial setup guarantees uniformity across sites and avoids inconsistencies in data processing.
2. **Quality Control (QC) and Principal Component Analysis (PCA):** Before initiating the analysis, the central server communicates agreed-upon QC parameters to all clients. Each client then independently performs QC on its local genotype data (e.g., filtering SNPs based on call rate, minor allele frequency, and Hardy-Weinberg equilibrium). After individual QC steps, the central server aggregates the lists of SNPs passing QC across all sites and identifies the intersection. The final set of target SNPs is then defined for the federated analysis. To correct for population structure, a federated PCA algorithm can be employed to generate the principal components. These components are essential for controlling for population stratification in the mixed model analysis while maintaining privacy.
3. **Federated Association Testing:** The core of the federated learning framework is federated association testing, which involves the distributed fitting of null regression models and mixed models, as well as the execution of a federated Score test. The final stage involves federated association testing, which is model-specific. This iterative process between the clients and the central server involves: (1) distributed fitting of the null regression model (linear or logistic), (2) distributed fitting of the null mixed model (linear or logistic), and (3) execution of a federated Score test to evaluate the association between each target SNP and the phenotype. Upon completion, the central server aggregates and disseminates the final GWAS results to all participants.

### 2 Model Construction

In this study, we propose a distributed framework for conducting GWAS using both generalized linear mixed models (GLMM) and linear mixed models (LMM), adapted for federated learning. These models are chosen for their ability to account for both fixed effects (such as covariates) and random effects (such as genetic relationships between individuals), which are critical for analyzing complex genetic data. The key distinction between GLMM and LMM is their applicability to different types of phenotype data: GLMM is suitable for binary phenotypes, such as disease status, while LMM is designed for continuous phenotypes, such as body mass index (BMI).

Building on the federated GWAS framework, we designed two distinct algorithms to handle these different phenotypic data types: the distributed GLMM for binary phenotypes and the distributed LMM for continuous phenotypes. Both algorithms leverage federated learning to ensure data privacy and facilitate efficient multi-center collaboration, while addressing the specific statistical needs of each model.

The federated generalized linear mixed model (FedGLMM) algorithm is specifically developed for analyzing binary phenotypes across multiple institutions without the need to share individual-level data. When using a GLMM with a logit link function for binary data, the model becomes a logistic mixed-effects model. Suppose there are *J* clients and one central server. The sample size for the *j* -th client is denoted as *n*_*j*_, and the total sample size across all clients is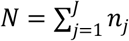. The binary phenotype of the *i*-th sample in the *j*-th client is represented by *y*_*ij*_ ∈ {0,1} .

For testing a single SNP, the algorithm considers the following distributed logistic mixed-effects model:

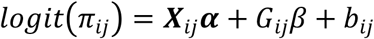

Here, *π*_*ij*_ = *P* (*y*_*ij*_ = 1|***X***_*ij*_, ***G***_*ij*_, *b*_*ij*_) is the probability that the binary phenotype of the *i*-th sample in the *j*-th client equals 1, given the covariates ***X***_*ij*_, the genotype ***G***_*ij*_, and the random effect *b*_*ij*_ . ***X***_*ij*_ represents the covariates for the *i*-th sample in the *j*-th client. ***G***_*ij*_ denotes the allele count (***G***_*ij*_ ∈ {0,1,2}) for a specific genetic variant (SNP) of the *i*-th sample in the *j*-th client. ***α*** is the fixed effects coefficient vector, and *β* represents the effect size of the genotype.

On a per-site basis, 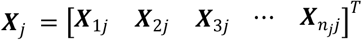 is the covariate matrix for the *j* -th client. The random effects vector for the *j* -th client,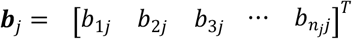, quantifies the random polygenic effects arising from cryptic relatedness. This vector is assumed to be sampled from a multivariate distribution with a mean of zero and a covariance matrix proportional to the kinship matrix _*j*_ . The global random effects vector is *b* = [***b***_1_ ***b***_2_ ***b***_3_ … ***b***_*J*_] ^*T*^, and it is assumed that *b* ∼ *N* {0, *τ* ***ξ***_***G*** *lobal*_, where *τ* is the global variance component parameter and ***ξ***_*Global*_ is the global kinship matrix for all samples across clients, which is an *N* × *N* matrix. When all individuals in the sample population are unrelated (i.e., there are no familial associations), the kinship matrices for all clients are identity matrices. The conditional expectation is *E* (*y*_*ij*_|*b*_*j*_) = *π*_*ij*_, and the conditional variance is *Var* (*y*_*ij*_|*b*_*j*_) = *ϕ ν* (*π*_*ij*_) = *ϕ π*_*ij*_ (1 − *π*_*ij*_ ). For binary data, the dispersion parameter *ϕ* = 1 . This model remains consistent for all genetic variants (SNPs).

In the linear mixed model, the dependent variable is assumed to follow a normal distribution and is a linear combination of the independent variables. There are a total of *J* clients and one central server. The sample size for the *j*-th client is denoted as *n*_*j*_, and the total sample size across all clients is 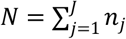. For testing a single genetic variant (SNP), the algorithm considers the following distributed linear mixed model:

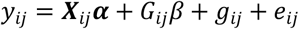

Here, *y*_*ij*_ represents the mean-centered continuous phenotypic value of the i-th sample in the *j*-th client. ***X***_*ij*_ denotes the covariate vector for the *i*-th sample in the *j*-th client. ***G***_*ij*_ represents the mean-centered allele count (taking values 0, 1,or 2) for the specific genetic variant (SNP) of the *i*-th sample in the *j*-th client. ***α*** is the fixed effect coefficient, and *β* denotes the effect size of the genotype. *g*_*ij*_ is the polygenic effect captured by kinship correlation, assumed to follow a normal distribution with a mean of 0 and a variance of *τ*_*g*_ ***ξ***_*j*_, where *τ*_*g*_ is the variance component parameter, and ***ξ***_*j*_ represents the kinship matrix among samples within the *j*-th client, which is an *n*_*j*_ × *n*_*j*_ -dimensional matrix. When the study population consists entirely of unrelated individuals (i.e, no familial relationships exist), the kinship matrix ***ξ***_*j*_ for all clients is the identity matrix. *e*_*ij*_ represents the random residual effect, assumed to follow a normal distribution with a mean of 0 and a variance of *τ*_*e*_***I***, where *τ*_*e*_ is the variance component parameter and ***I*** is the identity matrix.

At the site level, 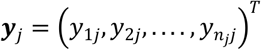 is the phenotypic vector for the *j* -th client. 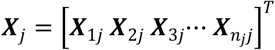 is the covariate matrix for the *j* -th client. The random effect vector for the *j* -th client, ***b***_*j*_ = [*b*_1*j*_*b*_2*j*_*b*_3*j*_…*b*_*n,j*_ ]^*T*^, quantifies the random polygenic effect due to cryptic relatedness. This vector is assumed to be sampled from a multivariate distribution with a mean of zero, and its covariance matrix is proportional to the kinship matrix ***ξ***_*j*_. The variance-covariance matrix for the phenotype ***y*** is ***V*** = *τ*_*g*_ ***ξ***_*Global*_ + *τ*_*e*_***I***. When the study population consists entirely of unrelated individuals (i.e, no familial relationships exist), the kinship matrices ***ξ***_*j*_ for all clients are identity matrices. In this scenario, *τ*_*g*_ and *τ*_*e*_ are not distinguishable. A single variance component parameter *τ*_*b*_ is considered instead of separately estimating *τ*_*g*_ and *τ*_*e*_ . Here, the combined random term is denoted as *b*_*ij*_ = *g*_*ij*_ + *e*_*ij*_, assumed to follow a normal distribution with a mean of 0 and a variance of *τ*_*b*_***I***. The linear mixed model can then be updated to a multiple linear regression model: *y*_*ij*_ = ***X***_*ij*_***α*** + ***G***_*ij*_*β*+ *b*_*ij*_,where the variance component parameter *τ*_*b*_ = *τ*_*g*_ + *τ*_*e*_ .

### 3 Algorithm Design and Optimization

The federated GWAS framework supports both binary and continuous phenotypes through two algorithmic implementations—FedGLMM and FedLMM—which share a three-stage optimization strategy. In the first stage, fixed effects are initialized via distributed generalized regression. In the second stage, model parameters are iteratively optimized using distributed PQL and AI-REML for generalized linear mixed models, and distributed AI-REML for linear mixed models. The final stage introduces a distributed GRAMMAR-Gamma approximation that enables efficient score testing across sites using parallel computation. Below, we present a detailed description of the three-stage optimization strategy used in FedGLMM.

#### Step 1: Initialization via Distributed Logistic Regression

For logistic mixed-effects models, the algorithm first obtains initial fixed-effects estimates by fitting a distributed logistic regression model, solved via a distributed Newton’s method. This strategy provides a superior starting point, leading to faster convergence in the subsequent mixed-model optimization than naive initialization.

The central server initiates the process by assigning initial values ***α***_0_ to the fixed-effects coefficients and defining a convergence threshold. These parameters are then transmitted to all participating clients. Each client proceeds to compute the local gradient 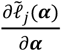 and Hessian matrix ***H***_*j*_ based on its covariate data and the received initial values. These local computations are subsequently sent back to the central server. The server aggregates the gradients and Hessian matrices from all clients to obtain the global gradient and global Hessian matrix. Using these global aggregates, the server updates the fixed-effects coefficients. This iterative process continues until the convergence criterion is met.

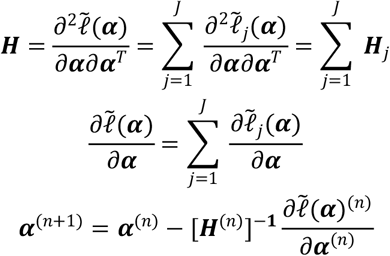

#### Step 2: Distributed logistic mixed-effects model

In the second step, the null distributed logistic mixed-effects model is fitted. The model parameters, including the fixed-effects coefficients, random-effects coefficients, and variance components, are estimated through an iterative optimization process that combines the distributed PQL method with the AI-REML algorithm.

Following initialization by the central server, which broadcasts starting values for the random effects *b*_0_, variance components ***τ***_0_ and the fixed-effects coefficients ***α***_0_ (from step 1), all clients engage in a multi-round iterative procedure with the server. This collaborative process implements the distributed PQL method combined with the distributed AI-REML algorithm to refine the parameter estimates.

Each client computes its local statistics based on its local data and then transmits these local values to the central server.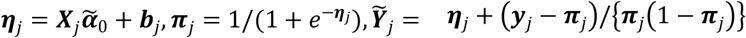. The server subsequently aggregates these local statistics across all clients to obtain the global values.

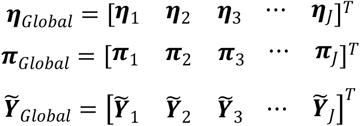

The global matrix ***W***_*Global*_ and ***Σ***_*Global*_ can then be calculated by the central server based on the global variables.

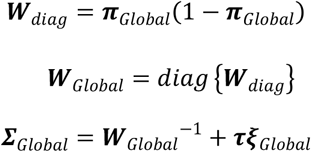

A critical component is the global genetic relationship matrix ***ξ***_*Global*_ . For an unrelated study population, this matrix is defined as an identity matrix of the corresponding dimension *N* × *N* . In studies with related individuals, a federated method for estimating the genetic relationship matrix must be employed. Wang et al.^33^ proposed SIGFRIED, a projection-based technique that leverages reference genotype data to estimate individual ancestry fractions, which are in turn used to infer relatedness in admixed populations within a privacy-preserving, federated framework. Alternatively, Wang et al.^34^ introduced SIGFRIED, a privacy-preserving and federated framework for estimating genetic kinship. The method is designed to accurately and efficiently compute kinship coefficients even for individuals from admixed populations.

Assuming ***ξ***_*Global*_ is a block-diagonal matrix, then the matrix ***Σ***_*Global*_ derived from it remains block-diagonal. The central server performs further calculations based on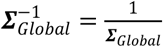. This matrix is then partitioned according to the sample size of each client, yielding a client-specific sub-matrix 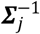 for each client, which is subsequently sent to the corresponding client. Upon receiving 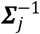 each client uses its local covariate data to compute its local intermediate results (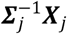 and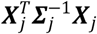). These local results are then transmitted back to the central server, which aggregates them to obtain the global variables.

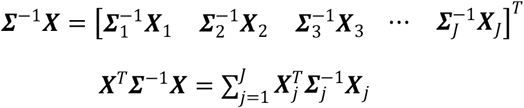

The central server then calculates the updated fixed-effects coefficients ***α*** and random effects *b*, using the existing global variables 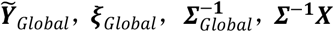 and ***X***^***T***^ ***Σ*** ^−1^***X*** from the current iteration.

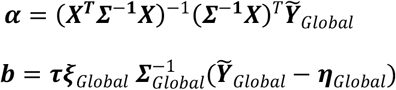

The central server then calculates the global projection matrix 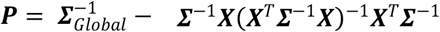 for that iteration’s step.

By defining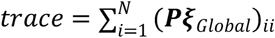, the term for the current iteration is derived.

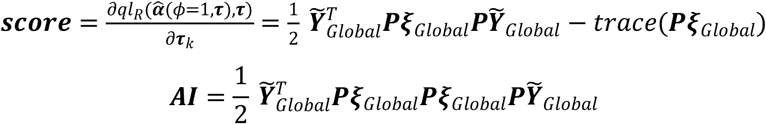

For the logistic mixed-effects model, *ϕ* = 1, ***θ*** = ***τ*** . Subsequently, an iterative process is employed within the REML framework. In each iteration, the distributed PQL method is used to obtain global vectors ***α***, working vectors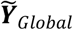, the projection matrix ***P***, and the information matrix ***AI*** . This iterative process, which updates alongside the parameters, continues until the optimal solution for the variance components ***τ*** is obtained. The specific steps of the iteration are as follows:

For each iteration *i*, the following steps are executed sequentially:

1. The variance component ***τ*** is updated preliminarily using the formula 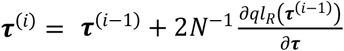
2. The global working vector 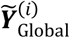 is updated based on the previous fixed effects estimates 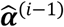 and the new *τ* ^(*i*)^ .
3. The Average Information matrix AI^(*i*)^ is computed.
4. The estimate of τ is refined using the inverse of the AI matrix: 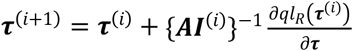.
5. The fixed effects 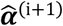 and random effects 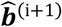 are updated utilizing the current global working vector 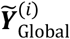 and the refined variance component *τ* ^(*i*+1)^ .
6. Finally, the global working vector is recalculated as 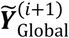 using the latest estimates of the effects.
7. This iterative cycle continues until a convergence criterion is met. The model is considered to have converged when the relative change in the parameter estimates between iterations falls below a predefined threshold *ϵ*, as defined by the condition:

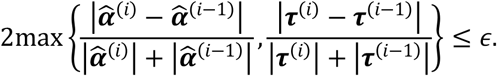

#### Step 3: Distributed Score test

Following the estimation of the mixed model parameters, a distributed score test is conducted to assess the association of each SNP while ensuring genotype data remains local. First, the central server partitions the global projection matrix ***P*** vertically according to each client’s sample size *n*_*j*_, which is 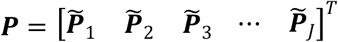, producing client-specific segments 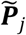. Each client receives its respective segment. It is important to note that 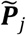 is an intermediate product of this distributed computation and not the actual client-specific projection matrix. Each client then calculates the local residual vector ***res***_*j*_ = ***y***_*j*_ − ***π***_*j*_ using its local phenotypic data ***y***_*j*_ and the predicted probability ***π***_*j*_ . Subsequently, leveraging its local genotype data ***G***_*j*_, each client computes the local statistics 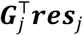 and 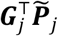, which are transmitted to the central server. The server aggregates these local statistics to obtain the global score statistic 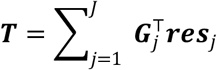 and the global matrix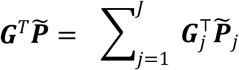. The server then performs a horizontal split on 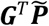 to generate client-specific segments 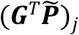 and sends them to the corresponding clients. Finally, each client computes local matrix products 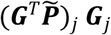 based on its genotype data ***G***_*j*_, which are aggregated by the server to obtain the final matrix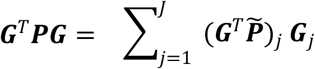. The variance of the score statistic ***T*** for each SNP is then obtained as *Var* (***T***) = *diag*(***G***^*T*^***PG***). Finally, the P-value for each SNP is calculated.

When sample sizes and the number of SNPs are very large, the aforementioned distributed strategy for calculating the test statistic and its variance requires repeated high-dimensional matrix operations. The resulting memory bottlenecks and prohibitive computational costs pose a critical obstacle to large-scale collaborative GWAS. To address this bottleneck, we propose a distributed GRAMMAR-Gamma approximation algorithm, which, combined with parallel computing techniques, achieves a significant improvement in computational efficiency. It is important to note that the GRAMMAR-Gamma approximation was originally proposed by Svishcheva et al.^35^ for linear mixed models. Subsequently, Zhou et al.^36^ in their work on the SAIGE method, demonstrated its applicability to the logistic mixed model scenario. The distributed GRAMMAR-Gamma approximation method introduced in this section is an extension built upon the foundation established for logistic mixed models in the SAIGE method.

To address the computational bottleneck of repeatedly calculating the variance-covariance matrix ***G***^***T***^***PG*** for a large number of SNPs in large-scale studies, we implemented a distributed version of the GRAMMAR-Gamma approximation. This method, based on a variance component model, employs a two-step procedure to significantly reduce computational cost.

The first step estimates a scaling factor *ŝ*. This is achieved by calculating the ratio 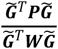 for a random subset of *m* SNPs (e.g., *m* =30), where 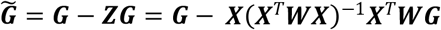 represents the covariate-adjusted genotype vector. The scaling factor *ŝ* is the mean of these ratios: 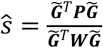. In the second step, the variance for the test statistic of all SNPs is efficiently approximated as: 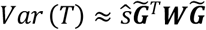, thereby circumventing the direct and costly computation of ***G***^***T***^***PG***.

Under data privacy protection requirements, which prohibit the transmission of any raw data, this study implements the GRAMMAR-Gamma approximation method within a distributed environment. The specific implementation strategy is as follows.

Since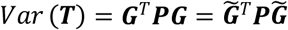, where 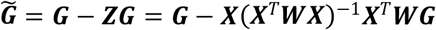, it is necessary to first compute the projection matrix ***Z*** = ***X*** (***X***^***T***^***WX***) ^−1^***X***^*T*^***W*** . The computation of ***Z*** proceeds in a distributed manner as follows. Given that the global weight matrix ***W***_*Global*_ is diagonal, the central server splits its diagonal elements ***W***_*diag*_ into *J* segments according to the sample size of each client, i.e., ***W***_*diag*_ = (*W*_1_, *W*_2_, …, ***W***_*J*_). The server then transmits the respective segment *W*_*j*_ to each corresponding client *j* . All clients subsequently compute the local matrix products 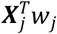 and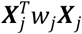, using their local covariate data ***X***_*j*_ . These local results are then sent to the first client (designated as *j*=1). Upon collecting all the local 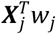 and 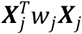 matrices, the first client aggregates them to obtain the global matrices 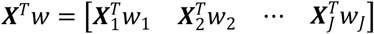 and 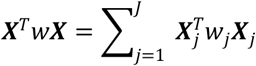. The first client then computes the inverse (***X***^***T***^ *w****X***)^−1^ and broadcasts this result to all other clients. Each client *j* then locally computes ***X***_*j*_(***X***^*T*^ *w****X***)^−1^ and sends the result back to the first client. The first client aggregates all the received ***X***_*j*_(***X***^*T*^ *w****X***)^−1^ matrices to obtain the global matrix ***X***(***X***^*T*^ *w****X***)^−1^ = [***X***_1_ (***X***^T^ *w****X***)^−1^ ***X***_2_ (***X***^*T*^ *w****X***)^−1^ ***X***_3_ (***X***^*T*^ *w****X***)^−1^ … ***X***_*J*_ (***X***^*T*^ *w****X***)^−1^]^*T*^ . The first client, now possessing the global matrices ***X***^*T*^*w* and ***X***(***X***^*T*^ *w****X***)^−1^, proceeds to compute the global projection matrix ***Z*** . Subsequently, the first client performs a horizontal split of ***Z*** into *J* segments, i.e., ***Z*** = [***Z***_1_ ***Z***_2_ … ***Z***_*J*_], and transmits each segment ***Z***_*j*_ to the corresponding client *j* . Upon receiving their respective segments ***Z***_*j*_, all clients compute the local matrix product ***Z***_*j*_***G***_*j*_ for all SNPs using their local genotype data. These local results are then sent to the central server. The server aggregates them to obtain the global matrix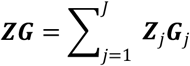. The central server then performs a vertical split on ***ZG***, resulting in segments ***ZG*** = [(***ZG***)_1_ (***ZG***)_2_ (***ZG***)_3_ … (***ZG***)] ^*T*^ . Each segment (***ZG***)_*j*_ is sent to the corresponding client *j* . Each client, upon receipt, calculates the covariate-adjusted genotype vector for each target SNP locally as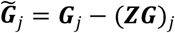. It is critical to note that 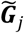 is a transformed intermediate result. This transformation makes it computationally infeasible to reverse-engineer the original genotype data ***G***_*j*_ from 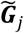, thus permitting its secure transmission. Clients then transmit their local 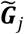 to the central server.

The central server aggregates these vectors to form the global adjusted genotype matrix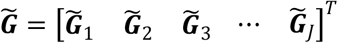. Using the existing global weight matrix ***W***_*Global*_, the server then computes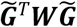. To implement the GRAMMAR-Gamma approximation, the central server directly extracts the genotype data 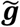 for a predefined set of *m* = 30 randomly selected SNPs from the global 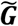 matrix. For each of these SNPs, the server calculates both 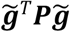 and 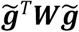. The scaling factor *ŝ* is then derived as the mean of the ratios of these quantities:

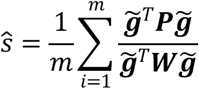

Finally, the variance of the score statistic for all SNPs is efficiently approximated as: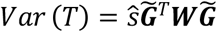. The entire federated score test procedure, including the distributed GRAMMAR-Gamma approximation, was implemented in C++. Computational efficiency was significantly enhanced by leveraging OpenMP multi-threading parallelization technology for multi-core processing.

### 3 Statistical Simulation Design

#### Genotype and Covariate Generation

Simulated datasets were derived from UK Biobank genotype data, restricted to unrelated individuals of British ancestry. In each simulation replicate, a random subset of samples and variants was selected to form the genotype matrix. Missing genotypes were imputed as random draws from a binomial distribution parameterized by the SNP-specific minor allele frequency (MAF). Covariates comprised sex (with a fixed effect of 0.1) and the top five principal components, the effects of which were randomly sampled from a standard normal distribution.

#### Phenotype Simulation

We generated phenotypes under a variance component model with a target narrow-sense heritability of *h*^2^ = 0.8. Twenty causal SNPs were randomly selected, with effects drawn from a standard normal distribution. We ensured orthogonality between genetic effects and covariates via projection, normalizing the total phenotypic variance to 1.

#### Experimental Design and Federated Setup

We assessed the performance of FedGLMM and FedLMM using a factorial simulation design varying in sample size (6,000, 15,000 samples) and feature dimension (50,000, 100,000, 200,000 SNPs). To mimic a realistic federated network, data were randomly partitioned across three clients and one central server. This design produced 12 distinct experimental scenarios per method, and each scenario was run over 100 independent replicates.

### 4 Empirical Analysis-Application to UK Biobank Data

#### Cohort and federated partitioning

To demonstrate real-world applicability, we applied the proposed FedGLMM and FedLMM algorithms to data from the UK Biobank. This large-scale cohort includes approximately 500,000 participants with comprehensive genetic and phenotypic profiles^15^. A subset of 100,000 unrelated British individuals was selected to ensure population homogeneity and high-quality genotypes. Participants were grouped into three federated sites based on geographic regions-Northern, Central, and Southern England.

#### Data processing and quality control

Genotype quality control was conducted using PLINK 1.9^37,38^ with standard thresholds (SNP missing rate < 0.02, sample missing rate < 0.02, MAF ≥ 0.01, Hardy-Weinberg Equilibrium P-value > 1×10-6, INFO ≥ 0.8). Samples with sex discrepancies or extreme heterozygosity were excluded. Each analysis adjusted for 15 covariates, including age, sex, age2, interaction terms (age2, sex × age, age2 × sex), and the top ten principal components computed via FlashPCA^39^. Two representative phenotypes were analyzed: smoking status (“ever” vs. “never”) for the binary case using FedGLMM, and body mass index (BMI) for the continuous case using FedLMM.

### 5 Evaluation Metrics

We employed a unified framework to assess both statistical accuracy and computational efficiency across simulation and empirical analyses. (1) **Statistical Accuracy**: To assess the calibration of test statistics, we generated Q–Q plots of score statistics, variance estimates, and P-values. The concordance of association discoveries was visualized using circular Manhattan plots, which directly quantify the overlap in significant loci between the federated algorithms and their centralized counterparts. We also computed the correlation of fixed-effects coefficients between methods for all analyses. In simulations with a known ground truth, statistical power and the false positive rate (FPR) were calculated to evaluate detection performance. (2) **Computational Efficiency**: For computational performance, three primary metrics were considered. Runtime encompassed the duration of the corresponding algorithm’s execution. Memory usage was measured as the memory consumption observed during the execution of the corresponding algorithm. Communication cost was quantified as the total volume of model parameters exchanged between clients and the central server throughout the federated training process, offering insight into the efficiency of the distributed computation framework.

### 6 Baselines

For benchmarking, we compared our federated implementations with fastGWA^9^ (continous) and fastGWA-GLMM^10^ (binary) from the GCTA^40^ toolkit. These centralized methods use mathematically equivalent mixed-model frameworks and are regarded as the “gold standard” for accuracy. We excluded two-step regression methods (e.g., REGENIE^11^) from the direct accuracy comparisons to ensure that the observed differences were due to the federated architecture rather than discrepancies in the statistical models. Comparative analyses were performed against established federated baselines, including FedGMMAT^29^. FedGMMAT was chosen as a federated reference due to its theoretical alignment with mixed-model frameworks. Methods based on two-step regression (e.g., SF-GWAS^27^) were excluded because of differences in statistical assumptions.

### 7 Experimental Setup

Our solution is deployed on a high-performance Linux server powered by the Montage Jintide® C6230R processor, boasting 96 cores and supported by an ample 500 GB of memory. The system runs on the stable and reliable Ubuntu 20.04.6 LTS operating system, providing an ideal environment for computational tasks. All analyses were conducted on this server, simulating a realistic federated network comprising three clients and one central server.

## Data Availability

Restrictions apply to the availability of these data. Data were obtained from the UK Biobank and are available at https://www.ukbiobank.ac.uk/ with the permission of the UK Biobank.

## Institutional Review Board Statement

This study was covered by the generic ethical approval for UK Biobank studies from the National Research Ethics Service Committee North West-Haydock (approval letter dated 17 June 2011, Ref 11/NW/0382).

## Informed Consent Statement

Informed consent was obtained from all subjects involved in the study.

## Acknowledgements

This research has been conducted using the UK Biobank Resource (Application #98273). All authors extend their sincere gratitude to all the participants and professionals contributing to the UK Biobank.

## Author Contributions

Xiaowen Suo: Conceptualization, Methodology, Software, Formal analysis, Investigation, Writing - Original Draft. Specifically, X.S. developed the FedLMM and FedGLMM algorithms, designed and implemented the federated learning framework, coded the software package, formulated the statistical models, conducted all simulation and empirical analyses, created the figures, interpreted the results, and wrote the first draft of the manuscript. Fuzhong Xue: Supervision, Project administration, Funding acquisition. Yanyan Zhao: Supervision, Project administration, Funding acquisition. All authors have read and approved the final manuscript.

